# Neural oscillations in the aging brain associated with interference control in word production

**DOI:** 10.1101/2022.08.26.505477

**Authors:** Xiaochen Y. Zheng, Vitória Piai

**Author notes:** Correspondence to Xiaochen Y. Zheng, Donders Centre for Cognitive Neuroimaging, Kapittelweg 29, 6525 EN, Nijmegen, The Netherlands.

## Abstract

Speaking is not only about retrieving words and structuring them into sentences, but it also requires top-down control to plan and execute speech. In previous electrophysiological research with young-adult speakers, mid-frontal theta oscillations have been observed using a picture-word interference paradigm. With this paradigm, participants name pictures while ignoring superimposed distractor words. In particular, mid-frontal theta power increases for categorically related distractors relative to other types of distractors, reflecting the top-down interference control in resolving the competition between processing streams during word production (Piai, Roelofs, Jensen, Schoffelen, & Bonnefond, 2014). In the present study, we conceptually replicated the magnetoencephalography study by Piai et al. (2014) with an older group of healthy adults (mean age of 60 years). Behaviorally, we replicated distractor semantic interference and Stroop-like interference effects usually observed in young adults. However, we did not find the corresponding theta modulation associated with these interference effects on the neural level. Instead, we found beta power decreases for both effects, mostly pronounced in the left posterior temporal and inferior parietal cortex. The distinct spectro-spatial-temporal profile of the oscillatory effects in the older population suggests different underlying dynamics relative to the midline frontal effect previously found in young-adult speakers. Our results indicate that the neural underpinnings of top-down interference control may be modified by aging, and that the mid-frontal theta cannot be the exclusive mechanism enabling interference control during spoken word production.

## Introduction

Speaking involves a set of processes that translate thoughts into words. It is not only about collecting the words and combining them into sentences or dialogues (Levelt, 1993; Levelt et al., 1999), but also requires sophisticated top-down control to plan and execute speech (Roelofs, 2021; Roelofs & Piai, 2011). In order to speak properly, one needs to plan and maintain the conversation goals and update the contents of working memory (Levelt et al., 1999; Martin & Slevc, 2014; Piai & Roelofs, 2013), to monitor what they have just said and what they are about to say (Hartsuiker, 2014), to choose between different words and to prevent interference from alternative words that get co-activated in the lexicon (Piai et al., 2013; Shao et al., 2015). Top-down control abilities, or in more general terms, *cognitive control*, change with advancing age, which impacts a wide range of cognitive domains, including working memory, attention, and inhibition (Braver & Barch, 2002; Paxton et al., 2007). Declines in the language domain, such as increased word-finding difficulties when speaking, are also commonly observed in older adults (Burke & Shafto, 2007; Schmitter-Edgecombe et al., 2000). However, it is still unclear how aging affects the cognitive control process required for speaking.

Cognitive control in language production is commonly investigated using the picture-word interference task (Hermans et al., 1998; Lupker, 1979; Piai et al., 2014; Shitova et al., 2017). In such a task, participants name pictures while ignoring simultaneously presented written or spoken distractor words. The distractor words may be, for example, semantically related (e.g., a picture of an apple combined with the distractor word “banana”) or unrelated (e.g., picture apple, distractor word “bike”) to the target picture name, or congruent with the target picture name (e.g., picture apple, distractor word “apple”). Naming performance, measured via response time (RT) or accuracy, changes as a function of the degree of relationship between the two competing streams (i.e., that of the picture and of the distractor). When the picture name and the distractor word are the same (i.e., congruent), their activations converge on a single word and further reduce the processing effort. By contrast, when the distractor word is incongruent with the target picture name, speakers need to inhibit the alternative word or enhance the target word (e.g., Piai, Roelofs, Jensen, et al., 2014; Roelofs, 2021; Shao et al., 2015) and prevent its interference, resulting in slower responses or more speech errors. Such a phenomenon is called *Stroop-like interference*, due to its shared nature with the classical Stroop effect (e.g., naming the red ink color of the printed word “blue”, Stroop, 1935). Conversely, when the distractor word is semantically related to the target picture name, it receives further activation from the picture and thus becomes a stronger competitor of the picture’s processing stream compared to a semantically unrelated distractor. Consequentially, the semantically related distractor causes larger delay and/or more errors than the unrelated ones, here referred to as *semantic interference*. This effect is well-established in the language production literature (Bürki et al., 2020; Glaser & Düngelhoff, 1984; Lupker, 1979). Both the semantic interference and the Stroop-like interference effects can be considered to reflect the top-down cognitive control employed to deal with distracting stimuli, sometimes also termed *interference control* (e.g., Friedman & Miyake, 2004; Friehs, Klaus, Singh, Frings, & Hartwigsen, 2020; Piai, Riès, & Swick, 2016).

Neuronal oscillations have been suggested to reflect the top-down interference control recruited for resolving the competition between processing streams during word production. In a magnetoencephalography (MEG) study, Piai, Roelofs, Jensen, et al. (2014) examined the neural mechanism underlying the competition between processing streams of the picture and the distractor word. A theta-power (4-8 Hz) increase was observed for the semantically related condition compared to the congruent condition (i.e., Stroop-like interference) roughly around 350-650 ms post-stimulus onset. The order of power increases was analogous to the behavioral effects (i.e., related > congruent). The theta effect was localized to the superior frontal gyrus and (left) postcentral gyrus, possibly also including the supplementary motor area and the anterior cingulate cortex (Figure 1, upper panel, modified from Piai, Roelofs, Jensen, et al., 2014).

**Figure 1.**
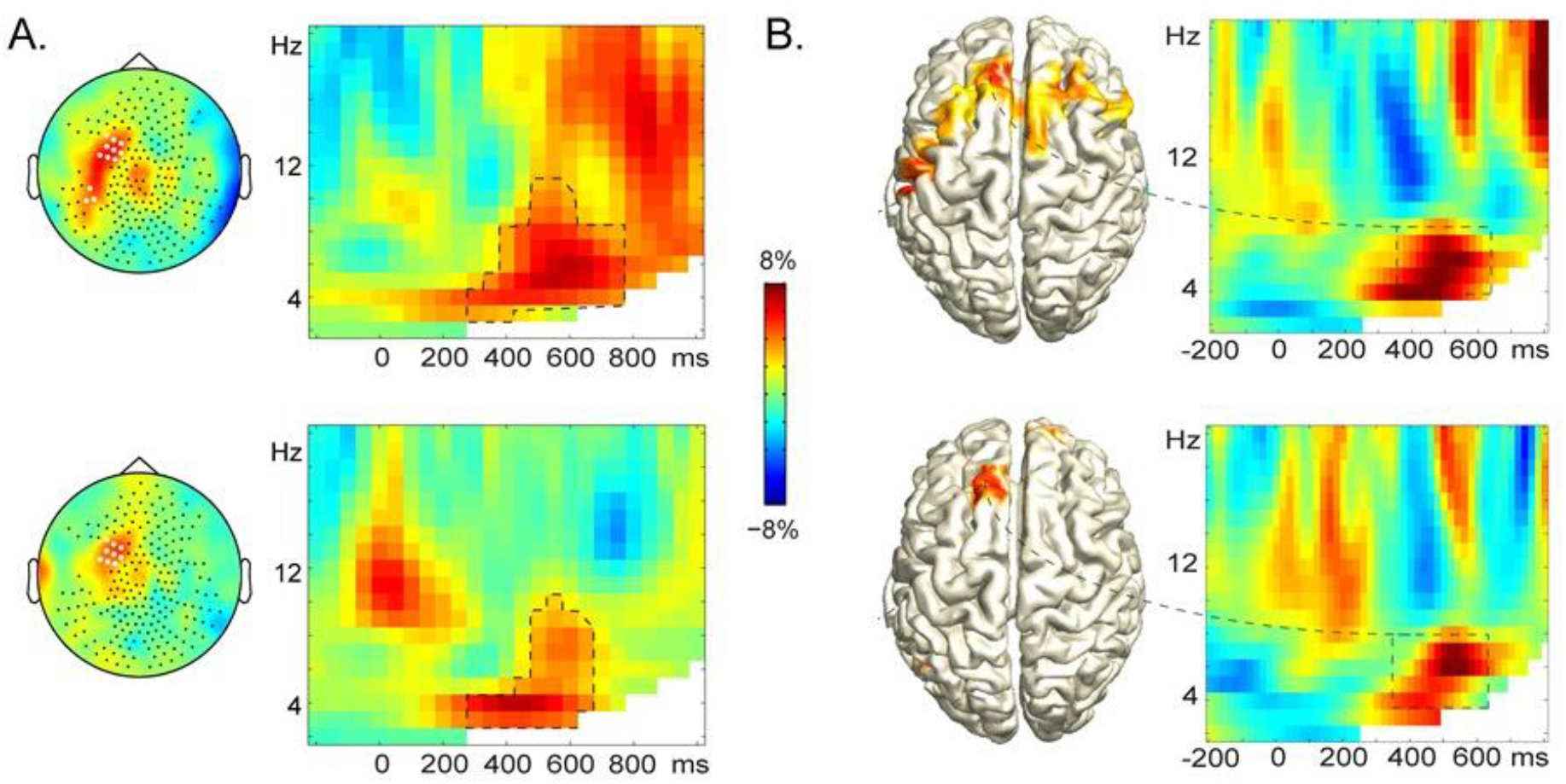
The mid-frontal theta power increase for Stroop-like interference (semantically related vs. congruent, upper panel) and semantic interference (semantically related vs. unrelated, lower panel). Color scale indicates the amount of relative power differences between the respective conditions. A. Sensor-level results: stimulus-locked time-frequency representations of relative power change, averaged over the sensors highlighted in white in the corresponding topographic maps to the left. B. Source-level results: estimated sources with their corresponding time-frequency representations. Figure modified from Piai, Roelofs, Jensen, et al., (2014). https://doi.org/10.1371/journal.pone.0088674.g001

Similarly, theta power increases in the same time window were also found for the semantic interference effect (i.e., related vs. unrelated), source localized again to the (left) superior frontal gyrus (Figure 1, lower panel). Both effects were interpreted as to reflect different degrees of effort, or top-down interference control, in resolving the competition among the competing stimuli. Similar mid-frontal theta power increases have also been observed in other language production studies, particularly in different conditions that require more control due to stimuli interfering with production processes (Krott, Medaglia, & Porcaro, 2019; Shitova et al., 2017; see Piai & Zheng, 2019 for a review on theta oscillation and cognitive control in language production). In the literature, midline frontal theta oscillations, generated by the anterior cingulate cortex and superior frontal gyrus, are associated with working-memory load (Itthipuripat et al., 2013; Jensen & Tesche, 2002), performance monitoring (Cavanagh et al., 2012; Cohen, 2011; Luu et al., 2004) and increased top-down control to prevent interference (Cohen et al., 2008; Cohen & Donner, 2013; Hanslmayr et al., 2008; Nigbur et al., 2011).

Piai, Roelofs, Jensen, et al. (2014) argued that mid-frontal theta oscillations would constitute the neural mechanism underlying top-down control needed for language production. However, their sample consisted only of young university students, whereas this conclusion might not extend to other groups, such as aging adults (Rad et al., 2018). For instance, language is a left lateralized function in the majority of the population but changes in lateralization may take place as compensatory or dedifferentiation processes in aging. The HAROLD (Hemispheric Asymmetry Reduction in Older Adults) model has proposed that the prefrontal activity during cognitive performances tends to be less lateralized in older adults compared to young adults (Cabeza, 2002, but see Berlingeri et al., 2013). Moreover, the right hemisphere involvement is strongly associated with cognitive reserve (Brosnan et al., 2018), which may also play a role in top-down control in healthy aging. The age-related changes can also occur in the form of frequency-band shift. Mid-frontal theta power has been found to be significantly lower in older than young adults in working memory tasks (Cummins & Finnigan, 2007; Kardos et al., 2014). Besides the changes in the theta dynamics with aging, the higher frequency bands (e.g., beta or gamma) become more apparent while the low frequency bands (e.g., delta or theta) diminish (Werkle-Bergner et al., 2006). How would neural oscillations, and in particular, the theta dynamics associated with the competition between processing streams in word production change for elderly adults? Could we safely draw the conclusion that mid-frontal theta reflects interference control for speaking across the life span?

In the present study, we perform a conceptual replication of Piai, Roelofs, Jensen, et al. (2014) with an older population, with the particular interest in how the mid-frontal theta is modulated by interference control, as measured through the Stroop-like interference and the semantic interference effects. We expect that the older population may show a different spectro-spatial-temporal profile of the cognitive control signature due to aging.

## Methods

### Participants

A recent study on word production has shown that age-related change in neurophysiological activity emerges from the age of 40 (Krethlow et al., 2021, in prep).With this in mind, we conducted the study with a group of participants around the age of 60 to have a more general insight on how a “no longer young” brain controls language production and prevents interference. Twenty-three participants from an older population (mean age = 61.0; range 46 -75, 13 males) than in Piai, Roelofs, Jensen, et al. (2014) took part in the study for monetary compensation. All of them were native speakers of Dutch, right-handed, with normal or correct-to normal vision.

The study was conducted in accordance with the Declaration of Helsinki, was approved by the local ethics committee (Arnhem-Nijmegen CMO, NL58437.091.17), and all participants provided written informed consent. Five participants’ data were excluded from the analyses due to excessive metal-related artifacts in the MEG and two participants were excluded due to too few correct responses, leaving a final sample of 16 participants (Mean age = 59.2, range = 46-72, nine males).

### Materials, Design and Behavioral Procedure

Eighty-eight color pictures were taken from the BOSS database (Brodeur et al., 2010), forming 16 different semantic categories with five to six objects pertaining to each category. For each picture, distractor words were the picture name (i.e., congruent), from the same semantic category as the picture (i.e., related), or from a different semantic category (i.e., unrelated, Figure 2). Thus, all distractor words belonged to the response set. We made sure that the related and unrelated distractors were phonologically unrelated to the target picture. All participants saw each picture once in each condition. The picture-word trials were randomized using Mix (van Casteren & Davis, 2006), with one unique list per participant. Participants saw no more than five consecutive trials from the same condition and there was no consecutive repetition of the distractor words or target pictures from trial to trial. Participants were instructed to name the picture and to ignore the distractor word. Both speed and accuracy were emphasized.

**Figure 2.**
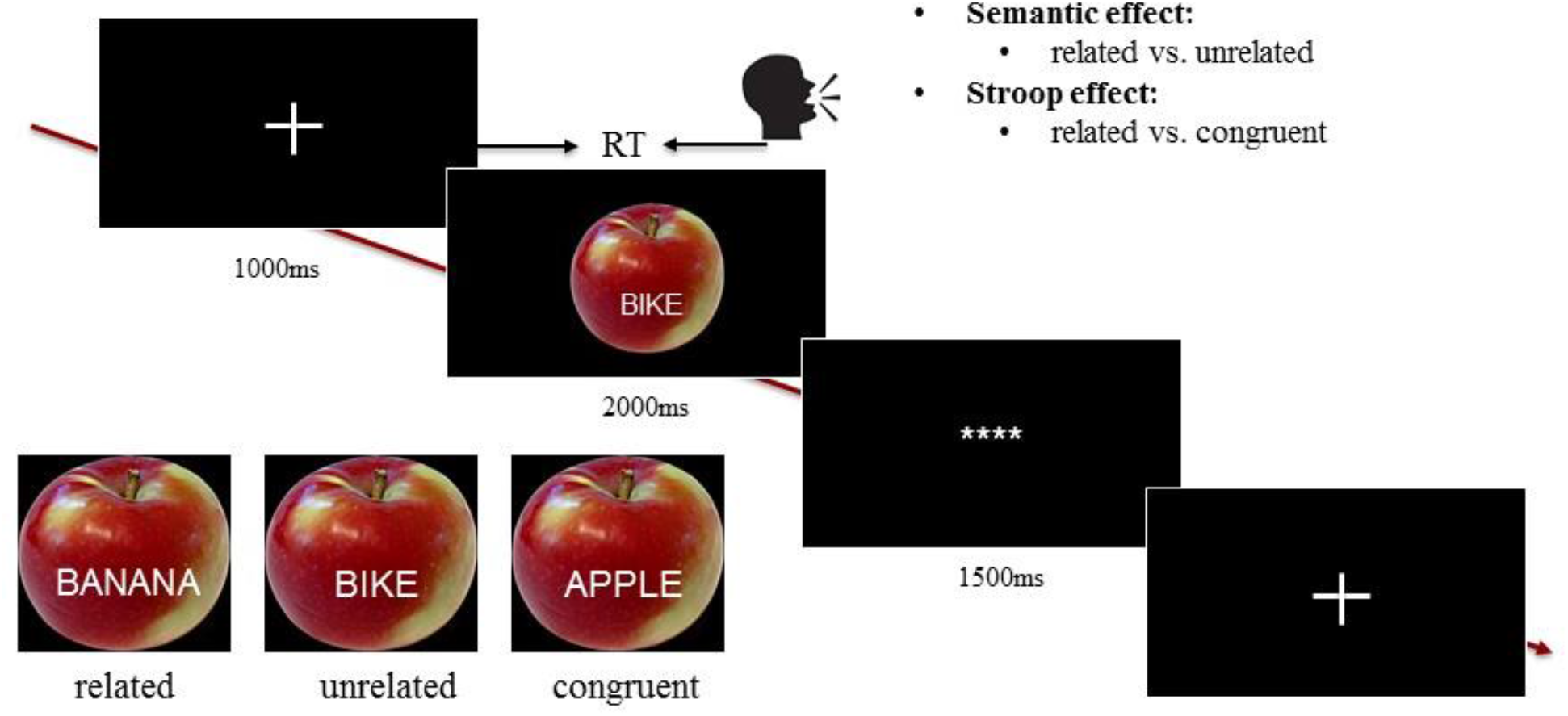
Experimental paradigm and conditions. The actual experimental stimuli are in the native language of participants.

The experiment was run using the software Presentation (Version 18.0, Neurobehavioural System Inc, Berkeley, U.S.). The background color of the computer screen was set to black, with a resolution of 1920 × 1080 pixels, at a refresh rate of 60 Hz. Distractor words were presented in white, centered on the picture. Trials began with a fixation cross presented for 1 s, followed by the presentation of the picture-word stimulus for 2 s. Then an inter-trial interval of 1.5 s was presented with **** on the screen (Figure 2). Participants’ responses were coded online as correct (i.e., identical to the target picture name) or incorrect. The experiment started with six practice trials. Participants were familiarized with the pictures and their names during the MEG preparation.

### MEG Acquisition

MEG data were acquired with a 271 axial gradiometer system (CTF Systems Inc., VSM MedTech Ltd.). The signals were analog low-pass filtered at 300 Hz, and digitized at a sampling rate of 1200 Hz. Participants were positioned in the MEG chair with pillows as they preferred. Three localization coils were attached to the participants’ head (the nasion, and the left and right ear canal). Throughout the measurement, the head position was continuously monitored using custom software (Stolk et al., 2013). Participants’ head position was re-adjusted when needed to maintain the original position (max. 5mm away). Four bipolar Ag/AgCl electrode pairs were used to measure the horizontal and vertical electrooculograms, the mouth electromyogram, and the electrocardiogram. Electrode impedance was kept below 20 kOhm. The MEG session lasted approximately 1 hr including preparation time.

Structural T1-weighted MRI scans of participants’ heads were acquired from a 3T Siemens scanner, either following the MEG session or on a different day no more than four weeks apart.

### Behavioral Data Analysis

Picture naming responses were recorded time locked to the picture onset, and RTs were manually calculated offline using the speech analysis program Praat (Boersma & Weenink, 2016), blinded for conditions. Responses containing incorrect picture names or disfluencies were considered as speech errors, and excluded together with technical errors from the subsequent RT and MEG analyses.

The statistical analyses of the behavioral data were performed with linear mixed-effects models using the lme4 package (Version 1.1.25, Bates et al., 2015) and lmerTest package (Kuznetsova et al., 2017) in R (Version 4.0.2; R Core Team, 2017). RTs were log-transformed to reduce skewness and approach a normal distribution. We used a model of RTs as a function of condition (related vs. unrelated vs. congruent, with the related condition as a reference), with a random slope of condition for participants as well as random intercepts for participants and pictures. Analysis of speech errors was done using a generalized mixed-effects model (binomial family), with random intercepts for participants and pictures.

### MEG Data Analysis

#### Preprocessing

We performed all MEG analyses using the Fieldtrip open source Matlab toolbox (Oostenveld et al., 2011) and custom analysis scripts in Matlab v.9.5.0 (R2018b, The Math Works, Inc). We first segmented the continuous MEG and EEG data into epochs from 500 ms before to 1000 ms after the target picture onset. The data were baseline corrected using the 500 ms interval before picture onset, down-sampled to 600 Hz, and then low-pass filtered with a cut-off of 55 Hz. EOG artifacts (i.e., eye blinks and saccades) as well as unambiguous speech artifacts were removed using independent component analysis, facilitated by visual inspection of the EEG data. Trials with remaining artifacts and/or bad sensors were rejected through visual inspection of the MEG data. This artifact rejection procedure was done before trials were separated by condition.

#### Sensor-Level Analysis

Synthetic planar gradients were computed for subsequent time-frequency (TFR) analysis (Bastiaansen & Knösche, 2000). To prevent contamination of the signal with speech-related artefacts, only the trials with RTs longer than 700 ms were included (for both sensor- and source-level analyses). The TFRs were computed between 500 ms pre-to 1s post-stimulus onset, at frequencies from 2 to 30 Hz, with a sliding time window of three cycle’s length, advancing in steps of 10 ms and 1 Hz. Each time window was multiplied with a Hanning taper. TFRs were baseline corrected using the 500 ms interval before stimulus onset, based on the normalized difference between the signal of interest and its baseline (i.e., S-B/(S+B)).

#### Source-Level Analysis

To examine the source of the beta effects found in the sensor-level analysis (see Results), we performed source localization of the effect in two time windows of interest (i.e., 200 ms to 600 ms post picture onset for the semantic effect and 200 to 500 ms for the Stroop-like effect). For each target window of interest, we defined its corresponding baseline window as a window of the same length prior to the target picture onset, e.g., a baseline window from 300 ms prior to picture onset would be used for a time window of interest from 200 to 500 ms post picture onset. We applied a frequency-domain beamforming technique (Dynamic Imaging of Coherent Sources, DICS) to all the sensor data (Gross et al., 2001).

Individual volume conduction models were computed using a realistic single-shell model (Nolte, 2003). The required brain–skull boundary was obtained from the participant-specific T1-weighted anatomical images, aligned to the MEG CTF coordinate system. To normalize across participants, we warped the CTF-aligned MRI scans to the Montreal Neurological Institute (MNI) space template to obtain participant-specific source model grids, with 10 mm resolution. Using the volume conduction models, lead field matrices were computed for each grid point for individual participants. Foreshadowing the results, sensor-level effects were not found in the 4-8 Hz frequency range, but rather in the 16-30 Hz range. This range formed the basis for the source-level analysis. The cross-spectral density matrix was computed at a central frequency of 23 Hz with 7Hz smoothing, resulting in a frequency range of 16-30 Hz. We computed a common spatial filter for both conditions combined (respective to the specific contrast) together with their corresponding pre-stimulus baseline windows, and then applied the spatial filter to the Fourier transformed sensor-level data per condition to estimate source-level power for each grid point.

The acquired sources for the time windows of interest were baseline corrected using corresponding sources for the baseline windows, in the same way as the sensor-level baseline correction (i.e., S-B/(S+B)). For visualization, the source-level results were interpolated to a template anatomical MRI.

#### Statistical testing

The statistical analysis of the MEG data was run using a nonparametric cluster-based permutation test (Maris & Oostenveld, 2007). On the sensor level, we compared two contrasts of interest, i.e., semantic effect (related vs. unrelated) and Stroop-like effect (related vs. congruent) for the theta frequency (4-8 Hz, based on Piai, Roelofs, Jensen, et al., 2014), as well as a broader frequency band (4-30 Hz) for exploratory analysis. This test provides a cluster-based *p* value (family-wise error corrected) of adjacent time-points, sensors and frequencies that exhibit a similar difference across conditions. Based on previous findings (Piai et al., 2014), we constrained the analyses of the theta band effect to the time window of 350 to 650 ms post stimulus onset and to all frontal and central MEG sensors (i.e., sensors labelled ‘MLF*’, ‘MRF*’, ‘MLC*’, ‘MRC*’). For the broader band exploratory analysis, we used the entire time window (i.e., 0-700 ms post stimulus onset) and all the available MEG sensors. The permutation distribution was constructed by randomly partitioning the original data 1,000 times. We considered spectro-spatial-temporal clusters with their cluster-level statistic corresponding to a p-value smaller than 0.05 to be significant.

## Results

### Picture naming performance

Figure 3 (left panel) shows the participants’ median naming RTs for each experimental condition. Participants were slower in the related condition (*M* = 1097, *SD* = 118) than in the unrelated condition (*M* = 1052, *SD* = 105), showing a significant semantic effect (β = 0.04, *SE* = 0.01, *t* = 4.89, *p* < .001). They were also slower in the related compared to the congruent condition (*M* = 936, *SD* = 105), showing a significant Stroop-like effect (β = 0.17, *SE* = 0.02, *t* = 8.80, *p* < .001). Similar results were found for the error rate (Figure 3, right panel). Participants made more naming errors for related (*M* = 6.5%, *SD* = 6.1%) compared to the unrelated condition (*M* = 4.4%, *SD* = 5.8%), showing a significant semantic effect (β = 0.47, *SE* = 0.18, *z* = 2.65, *p* = .008). They also made more errors for related compared to the congruent condition (*M* = 0.9%, *SD* = 1.5%), showing a significant Stroop-like effect (β = 2.23, *SE* = 0.29, *z* = 7.64, *p* < .001).

**Figure 3.**
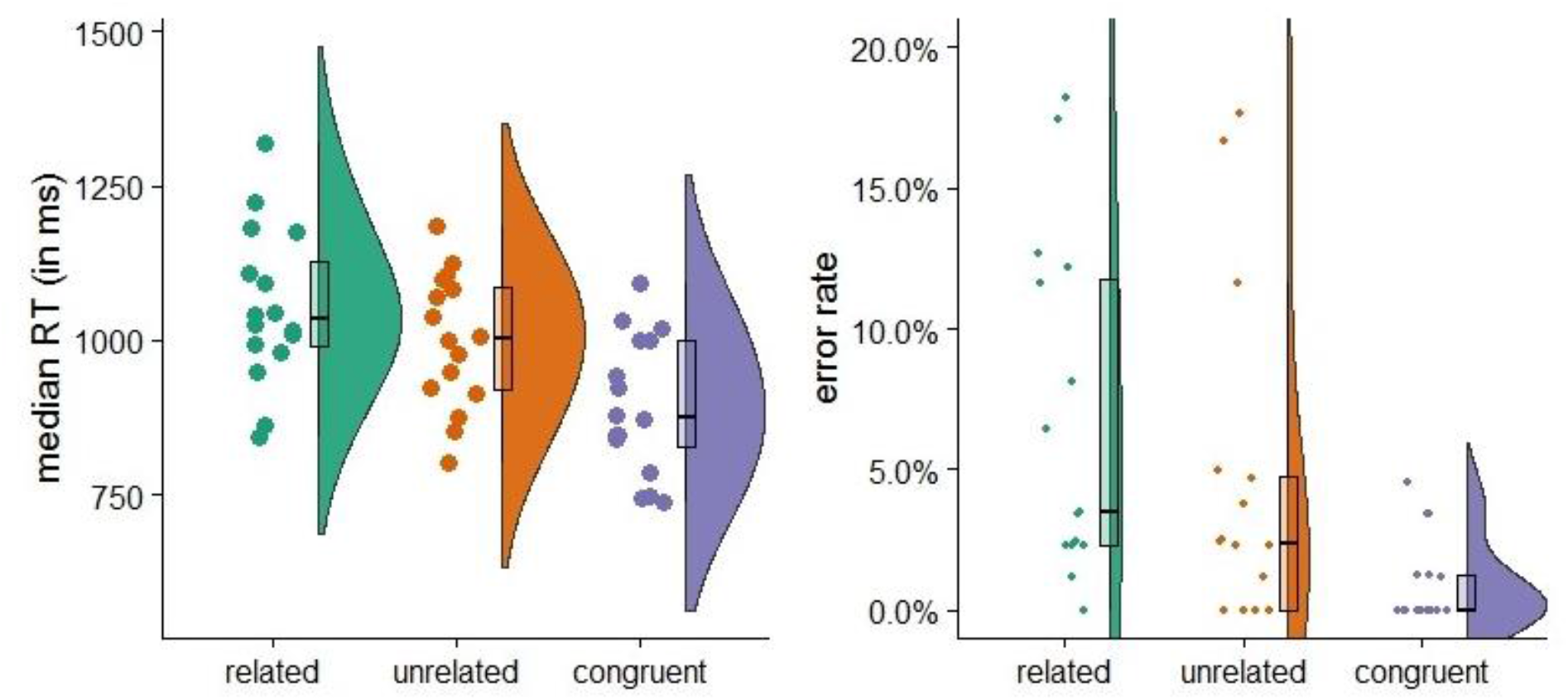
Raincloud plots of participants’ median response times (RTs, in ms, left panel) and error rates (right panel) for the three experimental conditions (related vs. unrelated vs. congruent). The outer shapes represent the distribution of the data over participants, the thick horizontal line inside the box indicates the group median, and the bottom and top of the box indicate the group-level first and third quartiles of each condition. Each dot represents one participant.

### Sensor-level effects

#### Semantic effect (related vs. unrelated)

The TFRs presented in Figure 4A show the power differences between related and unrelated conditions in the 4–30 Hz range between -500 and 700 ms time locked to the picture onset. The permutation test result shows that there is no significant difference between the related and unrelated conditions in the theta band (4-8 Hz) between 350 and 650 ms post picture onset for the fronto-central sensors (*p* = 0.506, Figure 4C). Nevertheless, visual inspection suggests a difference between the two conditions in the beta band. To explore the effect, we applied a cluster-based permutation test to all the sensors available in the full bandwidth (4-30 Hz) and full time window (0-700 ms). Results showed that there is a significant difference between the related and unrelated conditions in the beta band (*p* = .024). The effect is mostly pronounced around 200 to 600 ms post target picture onset, clustered around 16-30 Hz. The topographical map shows that the effect is mostly salient at the left central sensors (Figure 4B).

**Figure 4.**
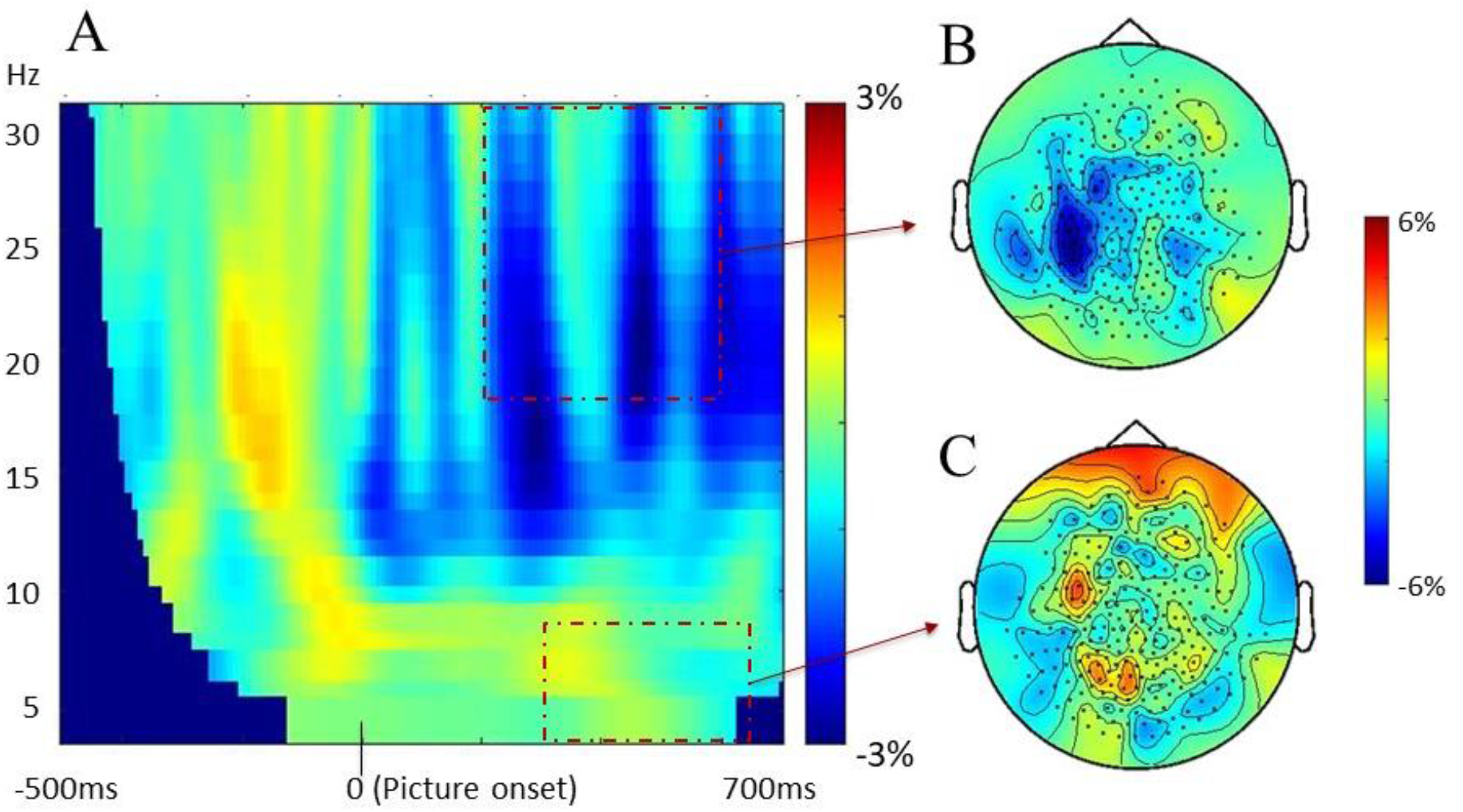
Semantic effect on the sensor level. (A) Stimulus-locked time-resolved spectrum of the contrast between related vs. unrelated conditions, averaged over all sensors. (B) Topography of the semantic contrast (i.e., related vs. unrelated) in the beta band (16-30 Hz) between 200 to 600 ms post picture onset (see the dashed line in 4A). (C) Topography of the semantic contrast in the theta band (4-8 Hz) between 350 to 650 ms post picture onset (see the dashed line in 4A). Color bar indicates the baseline-corrected power difference between conditions.

#### Stroop-like effect (related vs. congruent)

The TFRs presented in Figure 5A show the power differences between related and congruent conditions in the 4–30 Hz range between 0 and 700 ms time locked to the picture onset. The permutation test result showed that there is no significant difference between the related and congruent conditions in the theta band (4-8 Hz) between 350 and 650 ms post picture onset for the fronto-central channels (*p* = 0.677, Figure 5C). Similar to the semantic effect reported above, visual inspection again suggests a difference between the two conditions in the beta band. To explore the effect, we applied a permutation test to all the sensors in the full bandwidth (4-30 Hz) and full time window (0-700 ms). Contrary to what we found for semantic interference, the cluster-based permutation test for Stroop-like interference did not provide evidence in favor of the alternative hypothesis of differences between the related and congruent conditions in the beta band (*p* = .062). Nevertheless, given our a priori hypothesis of similar semantic and Stroop-like effects at the neural level and the fact that we established both semantic and Stroop-like effects at the behavioral level, we expect the lack of a statistically significant Stroop-like oscillation effect to be due to a power issue. To be able to perform source localization of the Stroop-like effect, also enabling comparability with the semantic effect, we applied a one-sided *t* test to inform on the spectro-temporal range of interest for running the subsequent source localization. This one-sided *t* test, which should be interpreted very cautiously, revealed an effect (*p* = .009) mostly pronounced between 200 to 500 ms post target picture onset, in the 16-30 Hz range, clustered around the central sensors (Figure 5B).

**Figure 5.**
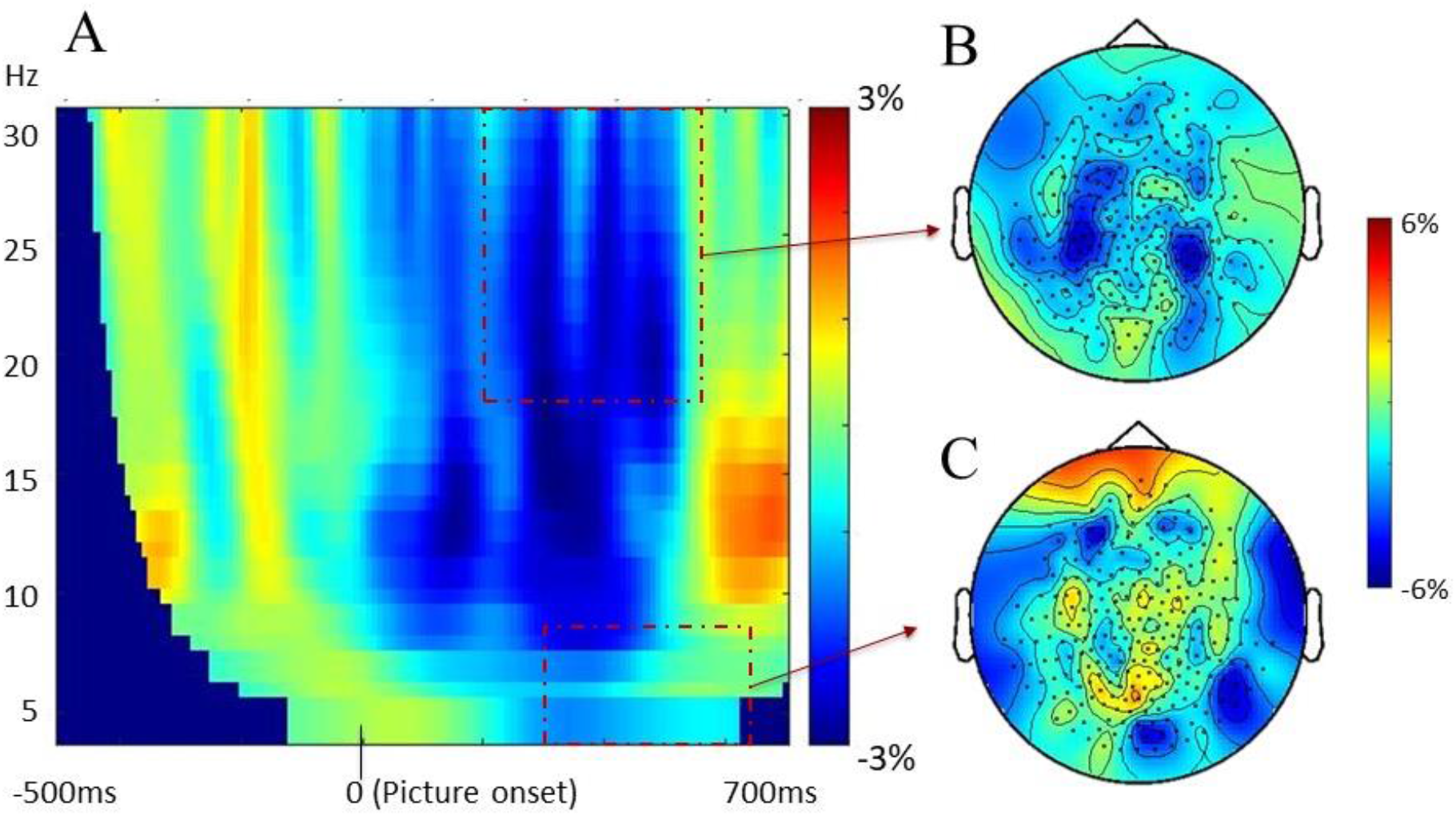
Stroop-like effect on the sensor level. (A) Stimuli-locked time-resolved spectrum of the contrast between related vs. congruent conditions, averaged over all sensors. (B) Topography of the Stroop-like contrast (i.e., related vs. congruent) in the beta band (16-30 Hz) between 200 to 500 ms post picture onset (see the dashed line in 5A). (C) Topography of the Stroop-like contrast in the theta band (4-8 Hz) between 350 to 650 ms post picture onset (see the dashed line in 5A). Color bar indicates the baseline-corrected power difference between conditions.

### Source localization of the semantic and the Stroop-like effects

To further understand the semantic and Stroop-like effects found on the sensor level, we performed source analyses in the beta band (16 to 30 Hz) for the semantic contrast (time window of 200 to 600 ms) and the Stroop-like contrast (200 to 500 ms). The results are shown in Figure 6. The beta power decreases associated with the semantic effect (*p* = 0.030) were found exclusively in the left hemisphere (Figure 6, top left), distributed around the pre- and post-central gyrus, supramarginal gyrus, and inferior parietal cortex, according to the AAL template (Tzourio-Mazoyer et al., 2002). The beta power decreases associated with the Stroop-like effect (*p* = 0.030) were also localized predominantly over the left hemispheres (Figure 6, bottom left), mostly posteriorly, including the angular gyrus, the middle temporal gyrus, and the inferior and superior parietal cortex. There were also some sources in frontal cortex, which were not visible for the semantic effect (Figure 6, right panels).

**Figure 6.**
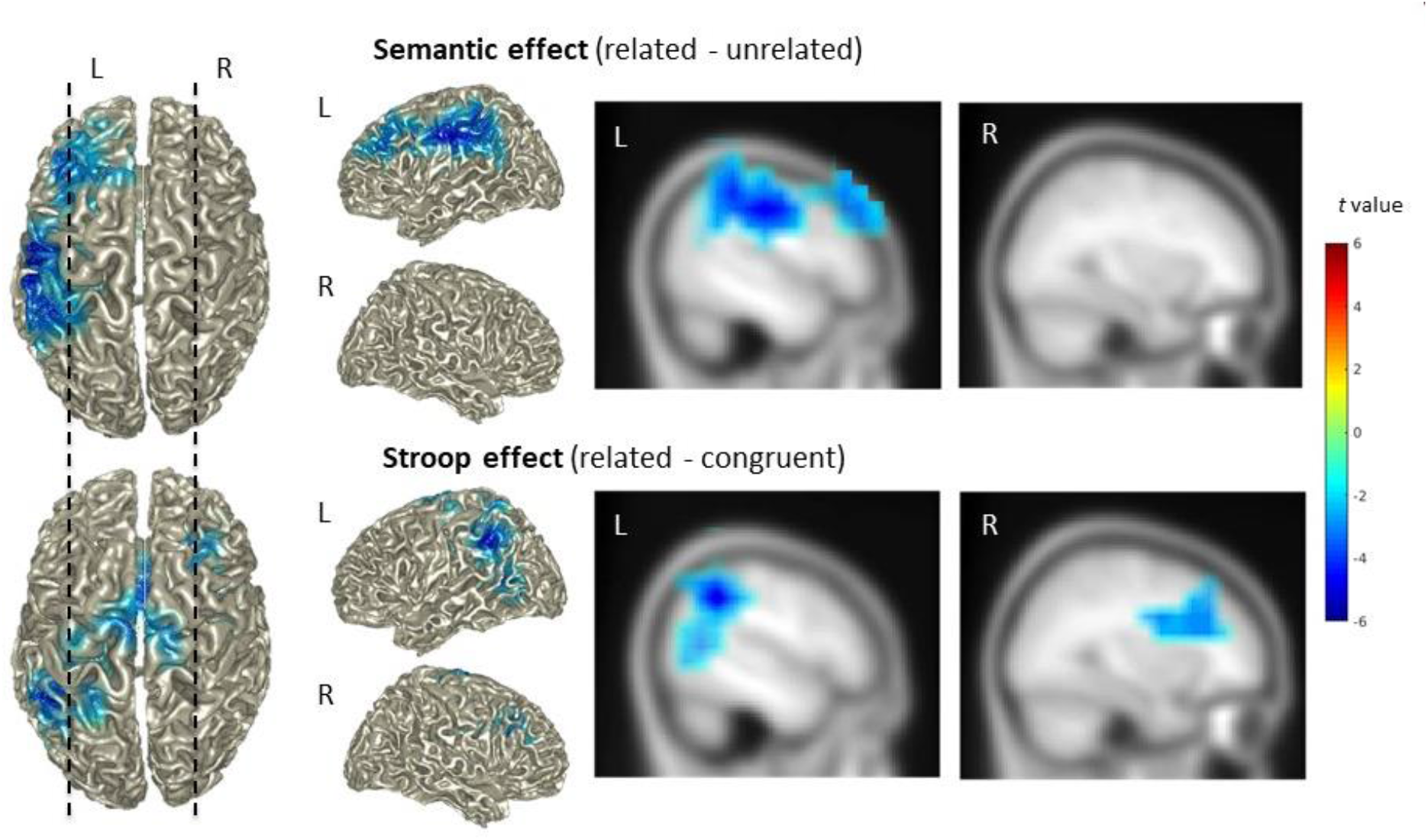
The semantic effect (top panel) and the Stroop-like effect (bottom panel) on the source level. Top left: Power difference between 16 and 30 Hz for the semantic effect (i.e., related vs. unrelated) in the time window of 200 to 600 ms post picture onset. Right: Two representative sagittal slices from the left and right hemisphere, corresponding to the dashed lines on the left. Bottom left: Power difference between 16 and 30 Hz for the Stroop-like effect (i.e., related vs. congruent) in the time window of 200 to 500 ms post picture onset. The two same sagittal slices for the Stroop-like effect are shown on the right. Color bar indicates the *t*-values for individual grid points. Only grid points associated with the significant cluster are colored.

## Discussion

In order to investigate age-related changes in interference control during spoken word production and associated changes in neuronal oscillations, we conceptually replicated the MEG study by Piai, Roelofs, Jensen, et al. (2014) in a group of aging adults with an average age of 60. Specifically, by using a picture-word interference task, we examined how the mid-frontal theta, as a marker for top-down cognitive control, is modulated by different types of interference. The behavioral pattern we found in the older adults was in line with what is commonly found in young adults: Speakers are better in naming pictures with congruent than with semantically related distractors, replicating the Stroop-like interference effect (Piai et al., 2014, 2020; Shitova et al., 2017); they are also better in naming pictures with semantically unrelated than with related distractors, replicating the semantic interference effect (Krott et al., 2019; Lupker, 1979; Piai et al., 2014, 2020; Schriefers et al., 1990). These results are also consistent with previous studies that examined picture-word interference in older populations (Graf et al., 1995; Rizio et al., 2017; Taylor & Burke, 2002). However, the neural signatures of the older adults that paralleled the two interference effects showed a very different profile than the young participants in Piai, Roelofs, Jensen, et al. (2014) (see Figure 1) and in other PWI studies (Krott et al., 2019; Shitova et al., 2017). Unlike the studies with young adults (mean age between 22 and 23), we did not find a mid-frontal theta effect either for the Stroop-like or the semantic effect. Instead, we found a significant beta power decrease accompanying both interference effects, which was not observed in the previous, young adults’ data in the original MEG study by Piai et al. Note though that the beta power decreases in the Stroop-like effect were not significant on the sensor level (*p* = .062, two-tailed test). Although slightly surprising, such dissociation between behavior and neural activity is in line with a recent finding by Krethlow et al. (2021), where age-related changes at the neurophysiological level starts from the age of 40, while the behavioral difference only appears after 70.

The theta power increases observed in picture-word interference tasks have been associated with top-down control in preventing the interference from the distractor words (Krott et al., 2019; Piai et al., 2014; Piai & Zheng, 2019; Shitova et al., 2017). The lack of a theta effect in this group of aging adults lends multiple interpretations. First – using a reverse-inference logic – it could suggest decreased abilities of detecting or suppressing interference in the older adults. It has been proposed that the attention-related prefrontal cortex function is altered with aging, leading to a decline in inhibitory control (Chao & Knight, 1997; West, 1996; but see e.g., Kramer, Humphrey, Larish, & Logan, 1994). For example, research has shown less theta power increase in older adults compared to younger adults during selective memory retrieval, suggesting the elderly being less capable of detecting memory interference (Ferreira et al., 2019). Nevertheless, the prolonged naming times instead of incorrect naming responses in most of the trials suggest that our older participants were still able to inhibit the irrelevant information and resolve the competition between the streams of processing the picture and the distractor. Therefore, the hypothesized deficit inhibition, if true, is limited in the neural level and not yet hampering performance on the behavioral level. Second, more recent theories of cognitive aging emphasize that the prefrontal cortex might not only be a major source of dysfunction but also a source of compensation (McDonough et al., 2013). An fMRI study on picture-word interference has reported that semantically related distractors led to increased recruitment of regions including the left middle frontal gyrus in older compared to the young adults (Rizio et al., 2017). It could be that our older adults instantly exhibit high-level (mid-frontal) theta activities for all the conditions due to (over-)compensation, resulting in a missing difference in the theta band between conditions. This assumption, however, is unlikely to be true in our data, given that the induced mid-frontal theta, as observed in the previous study of young adults, is not present in any of the three experimental conditions (cf. Figure 1, Figure S1). Third, it could be that the mid-frontal theta only reflects the top-down cognitive control of the young brain but not of the older brain, given that our older participants have clearly shown behavioral effects of interference control. Instead, interference control, at least in the older brain, can come about via other neuronal mechanisms than frontal theta oscillations.

Following the last account proposed above, is it possible that the beta power modulation we observed in the older adults is the same theta power modulation in the young adults (e.g., due to frequency band shifting)? This seems very unlikely. Besides the opposite polarity (i.e., power increase vs. decrease for the same effect), these neural signatures observed in our study and in Piai, Roelofs, Jensen, et al. (2014) are spatio-temporally distinct. While the theta increase is rather short lived and clustered around the 350-650 ms time window, the beta decrease is more widespread and starts relatively earlier in the processing. Besides, the interference-related theta power modulations have a fronto-central topography – therefore identified as the mid-frontal theta, a signature for cognitive control (Cavanagh et al., 2012; Cohen, 2011; Hanslmayr et al., 2008; Nigbur et al., 2011); whereas the present beta power modulations associated with the Stroop-like and semantic effects share a more left dominant, temporal-parietal profile – probably indicating a more language-related effect.

Therefore, another, perhaps more interesting, question is what the “new” beta can tell us about interference control during spoken word production. Before discussing its possible functionality, a first suspect we need to exclude is motor preparation. Motor-related activity is well-characterized by power decreases in the beta band in sensorimotor areas, typically in the range of 15-30 Hz (Cheyne, 2013; Pfurtscheller & Lopes da Silva, 1999). In language production, beta-band power decreases are observed prior to and during speech, in motor regions associated with speaking (Salmelin et al., 1995, 2000; Salmelin & Sams, 2002). Nevertheless, the beta power decreases we have observed are unlikely to be motor-related, or RT-related – as in that case, one would expect an earlier onset of beta power decreases for the congruent condition (i.e., shortest RTs) than the semantically related condition (i.e., longest RTs), resulting in a Stroop-like neural effect in the opposite direction of what we observed (i.e., beta power increase instead of decrease). The same reasoning holds for the semantic interference effect (i.e., RTrelated > RTunrelated). Alternatively, the beta effect could be related to working memory gating for task-relevant versus irrelevant information (Limanowski et al., 2020; Spitzer & Haegens, 2017), or semantic memory retrieval (Hanslmayr et al., 2009; Piai & Zheng, 2019). Alpha-beta power decreases in posterior temporal and inferior parietal brain areas have been reliably found in tasks that require conceptually driven word production, such as context-driven word production (Piai et al., 2015, 2018; Piai, Roelofs, & Maris, 2014), picture naming (Grappe et al., 2019), and verb generation (Pang et al., 2011). It is considered to reflect different degrees of retrieval of conceptual and lexical information from memory. The present results of beta power decreases resemble the spatial pattern found in context-driven word production (see e.g., Figure 5 in Roos & Piai, 2020). The similar recruitment of temporal-parietal brain regions may suggest their shared nature of semantic memory retrieval. However, this claim begs the question why the beta effect is not shown in the young population (e.g., see Figure 1). We do not have a good explanation for that. It could be that the beta power decreases are related to task demand. Given that retrieving is relatively easy for young adults in all three conditions, they may not show a difference in beta oscillations between conditions. However, at this point, this explanation is highly speculative.

The sources of power differences associated with distractor effects do not show a clear and complete overlap between the two types (i.e., semantic and Stroop-like), although they do show convergence in the left parietal lobe. The lack of a complete overlap at the source level between the two effects could be due to differences in signal-to-noise ratio at the level of the conditions compared, which would impact the source localization results. Additionally, although both effects tap into interference control, the conditions contrasted in each effect may also differ in other aspects, which could also affect which brain areas are recruited as part of a network.

In summary, with an older group of healthy adults, we replicated the behavioral Stroop-like interference and semantic interference effects commonly found in young adults, but not the corresponding theta modulation on the neural level (cf. Krott et al., 2019; Piai et al., 2014; Shitova et al., 2017). Instead, we found beta power decreases associated with both interference effects, mostly pronounced in the left posterior temporal and inferior parietal cortex. The distinct spectro-spatial-temporal profile of the oscillatory effects in the older population suggests different underlying dynamics other than the midline theta frontal effect. Our results indicate that, at the cognitive level, the top-down interference control process is present in aging adults, given the consistent behavioral effects we found. By contrast, its neural implementation seems to be different between the young adult and the old adult brains. Our findings suggest that the mid-frontal theta cannot be the exclusive mechanism enabling interference control during spoken word production.

## Acknowledgements

We thank Maaike ten Buuren and Rosemarije Weterings for help with data collection and behavioral data analysis.

## Data availability

Data are available at the Donders repository (https://doi.org/10.34973/6sa5-0t15).

## Supplementary material

**Figure S1.**
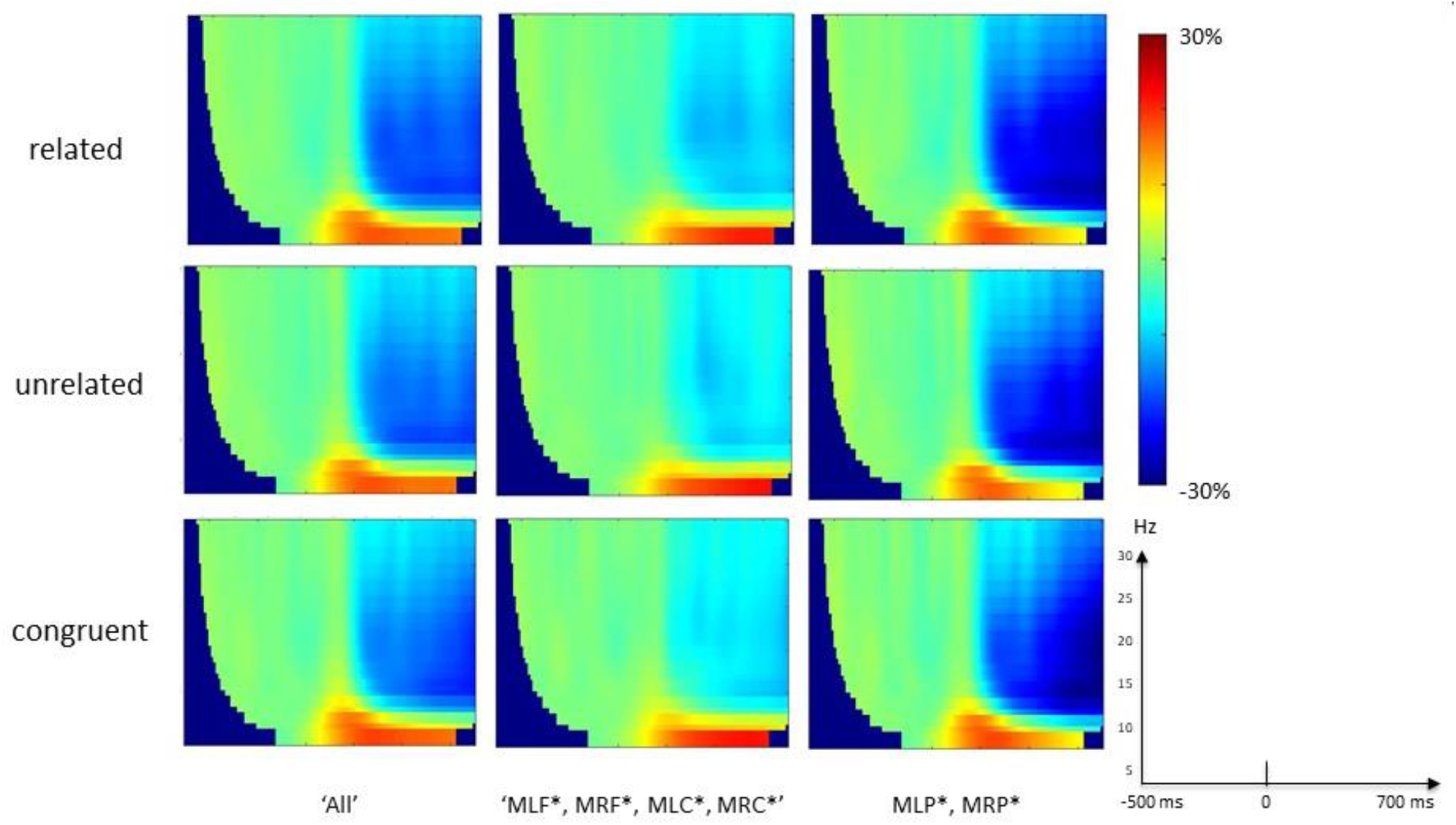
Time-frequency representation of the three experimental conditions, averaged across all sensors (left panel), fronto-central sensors (middle panel), or parietal sensors (right panel). All conditions are baseline corrected by the 500 ms time window before picture onset. Unlike Piai et al. (2014) (cf. Figure 1), data in the current study have only shown evoked theta power, regardless of experimental conditions and spatial profiles.

